# Sensory and motor electrophysiological mapping of the cerebellum in humans

**DOI:** 10.1101/2020.04.25.061044

**Authors:** Reiko Ashida, Peter Walsh, Jonathan C.W. Brooks, Richard J. Edwards, Nadia L. Cerminara, Richard Apps

**Author notes:** NLC, RJE and RA should be considered as joint senior authors. Corresponding author: Nadia Cerminara.

## Abstract

Damage to the cerebellum during posterior fossa surgery can lead to ataxia and in paediatric cases, the risk of cerebellar mutism syndrome. Animal electrophysiological and human imaging studies have shown compartmentalisation of sensorimotor and cognitive functions within the cerebellum. In the present study, electrophysiological monitoring of sensory and motor pathways was carried out to assess the location of limb sensorimotor representation within the human cerebellum, as a potential approach for real time assessment of neurophysiological integrity to reduce the incidence of cerebellar surgical morbidities.

Thirteen adult and paediatric patients undergoing posterior fossa surgery were recruited. For sensory mapping (n=8), electrical stimulation was applied to the median nerves, the posterior tibial nerves, or proximal and distal limb muscles and evoked field potential responses were sought on the cerebellar surface. For motor mapping (n=5), electrical stimulation was applied to the surface of the cerebellum and evoked EMG responses were sought in facial and limb muscles.

Evoked potentials on the cerebellar surface were found in two patients (25% of cases). In one patient, the evoked response was located on the surface of the right inferior posterior cerebellum in response to stimulation of the right leg. In the second patient, stimulation of the extensor digitorum muscle in the left forearm evoked a response on the surface of the left inferior posterior lobe. In the motor mapping cases no evoked EMG responses could be found.

Intraoperative electrophysiological mapping, therefore, indicates it is possible to record evoked potentials on the surface of the human cerebellum in response to peripheral stimulation.

## Introduction

The cerebellum is involved in the coordination of voluntary movements, postural balance and learning of new motor skills (1). However, an increasing body of evidence indicates that the role of the cerebellum also extends to cognitive functions (2–8). Surgical damage to the cerebellum results in ataxia, and in children, posterior fossa surgery can lead to cerebellar mutism syndrome in up to a third of patients, characterised by a transient loss of speech, behavioural impairments, emotional lability and hypotonia (9–11).

Extensive anatomical and electrophysiological mapping studies in non-human species have shown that the cerebellum and its associated input/output pathways are functionally compartmentalised into modules (12–14). In humans, magnetoencephalography (MEG) studies have reported short-latency responses (ca. 13-19 ms) in the cerebellum, evoked by median nerve stimulation (15, 16). As in other mammalian species, peripheral stimulation is therefore capable of synchronous activation of populations of neurons in the human cerebellum to generate substantial field potentials.

While non-invasive neurophysiological techniques have the temporal resolution to reveal such responses in humans, these techniques require considerable averaging to detect cerebellar responses. Direct electrophysiological recording from the cerebellum overcomes this problem. To date, two studies have explored this possibility. Preliminary studies by Mottolese et al., (17, 18) reported evoked potentials in the posterior cerebellum (lobule VI), in response to stimulation of the hand and mouth muscles, while Hurlbert et al., (19) also recorded evoked potentials from the posterior cerebellum in humans but in response to stimulation of the tibial nerve.

The present study extended these findings by exploring the possibility of recording field potentials from the surface of the human cerebellum evoked by upper or lower limb stimulation, as well as directly stimulating the cerebellar surface to determine if peripheral EMG responses can also be evoked. If either approach was successful, the findings could provide the basis for subsequent clinical application as a method to minimise damage to ‘eloquent’ cerebellar areas.

## Patients and Methods

### Patients

Study approval was obtained by the National Research Ethics Service Committee South West Service, Frenchay Hospital, North Bristol Trust (REC reference 12/SW/0050) and University Hospitals Bristol NHS Foundation Trust (UH Bristol CH/2014/4599). The study was conducted according to the Declaration of Helsinki 2013. Any patient undergoing a posterior fossa craniotomy over the age of two, fluent in English, total operation duration of more than three hours and those without any contraindications to neurophysiological monitoring were included in the study. Patients with previous history of posterior fossa craniotomy and with any pre-existing neurological conditions were excluded from recruitment.

Thirteen patients (nine male) were recruited in total. Age range was 3-63 years (median age 24 years). Eight patients underwent peripheral electrical stimulation and sensory mapping of the cerebellum. Five patients underwent cerebellar cortical electrical stimulation for motor mapping.

### Surgery

Surgery was performed under propofol anaesthesia to minimise interference with neurophysiological monitoring (20). Patients were positioned either prone or seated with their heads secured in Mayfield® skull clamps. A midline posterior fossa craniotomy approach was used for all patients. Patients were registered to a neuro navigation system by surface registration to the uploaded T1 fine cut axial MRI with gadolinium. The size of the craniotomy and cerebellar exposure varied between patients depending on the location of the pathology.

Standard intraoperative monitoring of motor and sensory evoked potentials (MEPs and SEPs) and cranial nerve activity allowed monitoring of the corticospinal tract and somatosensory pathways (e.g. dorsal column-medial leminiscal pathway) and the afferent nerve volley. EMG recordings using twisted pair needle electrodes (Ambu, Copenhagen, Denmark) were obtained from the muscles of the face, oropharynx and shoulders supplied by the cranial nerves, small muscles of the hands (abductor pollicis brevis, the adductor digiti minimi or the 1st dorsal interosseous) and abductor hallucis brevis (21). SEP recordings were obtained from scalp using corkscrew electrodes (Ambu Copenhage, Denmark) placed over the contralateral parietal lobe (21), in response to the stimulation of the median nerve (upper limb) and posterior tibial nerve (lower limb) (Ambu disposable Neuroline stick on electrodes). Cerebellar evoked potentials were recorded from the cerebellar cortex (see below for more detail). Corkscrew stimulation electrodes (Ambu, Copenhagen, Denmark) were positioned on the scalp overlying the precentral gyrus for monitoring MEPs in the small muscles of the hands and in abductor hallucis brevis.

### Limb stimulation

Peripheral stimuli were delivered by a 32 channel Neuromaster IOM system (Nihon Kohden, Tokyo, Japan). For the first patient (S1), single pulse constant current (0.2 ms duration) electrical stimulation at a rate of 5.1 Hz for the upper limb stimulation and 3.1 Hz for the lower limb stimulation were delivered, identical to the stimulation parameters used for standard clinical SEP monitoring in patients (22). Parameters were modified for subsequent cases (cases S2-8) based on those used in animal studies and stimulus rates ranged from 0.2-0.5 Hz (23, 24). In some cases, paired pulse stimulation was delivered (1 ms inter-stimulus interval). The peripheral stimulus intensity was adjusted to evoke a small but detectable twitch in the corresponding body part. This was approximately 20 mA for the arms and 30-40 mA for the legs.

Three patients (S6-8) underwent sensory mapping of the cerebellum using stimulus parameters based on the study performed by Mottolese et al., (18) (Table 1). Electrical trains of nine pulses, 0.5 ms duration, and an inter stimulus interval of 10 ms were delivered at 2.7 Hz at an intensity which produced a muscle twitch. Using these parameters, in one patient (S6) the forearm extensors and the tibialis anterior muscles in the lower leg were stimulated using twisted pair needle electrodes (Ambu, Copenhagen, Denmark), in addition to median and posterior tibial nerves. More proximal limb muscles, biceps (upper arm) and quadriceps (thigh) were stimulated for the two other patients (S7, S8). Table 1 indicates the different stimulus parameters and limb stimulation sites used for all eight sensory mapping patients.

**Table 1.** Table showing the summary of recruited patient demographic and stimulation parameters for sensory mapping

| Case no | Age | Sex | Histology | Location of pathology | Stimulation location | Peripheral stimulation parameters |
| --- | --- | --- | --- | --- | --- | --- |
| S1 | 47 | F | Ganglioglioma | Tectum<br>IVth ventricle | Median and posterior tibial nerves | Single 0.2 ms square pulse, 5.1 Hz, posterior tibial nerve<br>Single 0.2 ms square pulse 3.1 Hz, median nerve |
| S2 | 49 | M | Metastasis unknown primary | Cerebellar hemisphere | Median and posterior tibial nerves | Single 0.2 ms square pulse, 0.5 Hz |
| S3 | 7 | M | Ependymoma | IVth ventricle | Median and posterior tibial nerves | Single 0.2 ms square pulse, 0.5 Hz |
| S4 | 27 | M | Cavernoma | Cerebellar vermis | Median and posterior tibial nerves | Paired 0.2 ms square pulses, 1 ms interval at 0.5 Hz |
| S5 | 39 | M | Arachnoid cyst | Supra cerebellar | Median and posterior tibial nerves | Paired 0.2 ms square pulses, 1 ms interval at 0.5 Hz |
| S6 | 16 | F | Pilocytic astrocytoma | Cerebellar hemisphere | Lower arm, lower leg | 9 square pulses 0.5 ms, 10 ms interval, 2.7 Hz |
| S7 | 6 | M | Ependymoma | IVth ventricle | Upper arm, thigh | 9 square pulses 0.5 ms, 10 ms interval, 2.7 Hz |
| S8 | 24 | M | Diffuse astrocytoma | Brainstem | Upper arm and thigh | 9 square pulses 0.5 ms, 10 ms interval, 2.7 Hz |

### Sensory mapping: cerebellar evoked potentials

Electrophysiological recording from the exposed surface of the cerebellum (dura reflected) was carried out prior to tumour resection in the eight sensory mapping patients (Table 1). A bipolar 2mm ball tipped stimulation probe (Inomed, Emmendingen, Germany) with 5-8 mm width between the two contact points was used for seven of the eight patients (S1-4, S6-7). Electrophysiological signals were recorded differentially between one of the contact points and an indifferent electrode placed nearby in the subcutaneous tissue alongside the incision. The data were amplified (x1000) and bandpass filtered (30 Hz to 3 KHz). The probe was held free hand for the first patient (S1) and gently placed at different positions on the cerebellar cortical surface. For the remaining patients (S2-S7), the recording probe was fixed in a flexible arm retractor to minimise movement artefacts during recording. In all cases, the bipolar probe was moved in increments of approximately 5 mm laterally and rostro-caudally in a systematic manner to cover the entire exposed cerebellar surface. A four-contact recording strip with 10mm spacing (Ad-tech Medical Instrument Corporation, Oak Creek, USA) was used in one patient (S5). SEP recordings provided a positive control that the peripheral stimulation was effective in generating an ascending sensory volley (Figure 2).

### Motor mapping: cerebellar stimulation

In five patients (M1-5) the cortical surface of the cerebellum was stimulated using a monopolar probe in order to evoke EMG responses from the nasalis and orbicularis oris, biceps, forearm (extensor digitorum communis and flexor carpi radialis), small hand muscles, quadriceps, tibialis anterior and abductor hallucis, using parameters based on human intraoperative transcranial MEP monitoring and animal and human cerebellar stimulation (Table 2, (17, 25)).

**Table 2.** Table showing the summary of recruited patient demographic, stimulation parameters for motor mapping.

| Case no | Age | Sex | Histology | Location of pathology | Cerebellar stimulation parameters |
| --- | --- | --- | --- | --- | --- |
| M1 | 55 | M | Ependymoma | IVth ventricle | 5 pulses, pulse duration 0.3 ms, 400 Hz, 10, 20, 30 mA, 11.5ms total stim duration |
| M2 | 3 | F | Pineal blastoma | Pineal | 5 pulses, pulse duration 0.3 ms, 400 Hz, 10, 20, 30 mA, 11.5 ms total stim duration |
| M3 | 11 | M | Pineal blastoma | Pineal | 35 pulses, 150 Hz, pulse duration 0.3 ms, 10 mA, 235 ms total stim duration, |
| M4 | 63 | M | Ependymoma | IVth ventricle | 35 pulses, 150 Hz, pulse duration 0.3 ms, 10 mA 235 ms total stim duration, |
| M5 | 6 | F | Ependymoma | IVth ventricle | 35 pulses, 150 Hz, pulse duration 0.3 ms, 10 mA 235 ms total stim duration, |

A train of five anodal square wave pulses with an individual pulse duration of 0.3 ms and an inter-stimulus interval of 2.5 ms was delivered at each stimulation site in two patients (M1-2) with a stimulus strength of 10-30 mA. For the subsequent three patients (M3-5), a train of 35 anodal square pulses with an individual pulse duration of 0.3 ms and an inter-stimulus interval of 2.5 ms was delivered at each stimulation site with a stimulus intensity of 10 mA. Bipolar stimulation was also carried out in one patient (M3) using the same bipolar probe. Cathodal stimulation was carried out in one patient (M2, see Table 2). Charge density ranged from 0.7-4.5 micro C/cm2/phase. This was below the maximum safe charge density (up to 7.4 micro C/cm2/phase) based on human chronic cerebellar stimulation and non-human primate cerebellar stimulation (26, 27).

### Statistical and data analysis

Data were analysed offline using Spike2 software (CED, Cambridge, UK). Approximately 30 trials were averaged per recording site for six sensory mapping patients (S1-5, S8). For the two remaining sensory mapping patients (S6,7) 100 to 150 trials were averaged per recording site. Recordings from every cerebellar cortical recording site for the sensory mapping cases, and the EMG recording from peripheral muscles for the motor mapping were carefully examined for evoked potentials at the time of recording. If a response was evident, then average onset latency, which was taken from the first stimulation pulse, and peak to peak amplitude were measured offline.

### Functional MRI

In one motor mapping patient (63 year-old male, M4) with a fourth ventricular ependymoma, pre-operative functional MRI was undertaken. This was done in order to increase the likelihood of locating sites for cerebellar stimulation to evoke a peripheral response. The motor fMRI paradigm involved the patient moving their fingers or toes at an irregular rhythm directed by the flashing words ‘fingers’ or ‘toes’ on an LCD screen(8). fMRI data were analysed using the FSL software package (http://fsl.fmrib.ox.ac.uk). The functional data were uploaded onto the Stealth navigation system (Medtronic, Minneapolis, USA). The analysed fMRI data and the T1 structural scan (Magnetization-Prepared Rapid Gradient-Echo sequence, MPRAGE(28)) acquired at CRiC Bristol (University of Bristol) were in NIfTI (Neuroimaging Informatics Technology Initiative) format. These were converted to DICOM (Digital Imaging and Communications in Medicine) format which was compatible to the Stealth navigation system. The converted fMRI and structural images were uploaded to Stealth. These were then transformed onto the T1 structural MRI scan, which was acquired at the operating hospital, which was the MRI used for clinical navigation. Cerebellar surface cortical stimulation was carried out at the closest, accessible surface site from the BOLD activated area within the cerebellum.

## Results

### Sensory mapping

In eight patients, peripheral limb stimulation (cases S1-S8) was used to determine if evoked field responses could be recorded from the cerebellar surface during posterior fossa surgery. Figure 1a shows an example evoked potential recorded from the surface of the right inferior posterior lobe of the cerebellum with an onset latency of 13 ms in response to stimulation of the ipsilateral posterior tibial nerve. Evoked potentials of a similar peak-to-peak amplitude (~16 μV) and onset latency were recorded at six adjacent recording positions on the cerebellar surface. By contrast, stimulation of the ipsilateral right arm at the same recording sites failed to evoke a detectable response (Fig. 1a).

**Figure 1.**
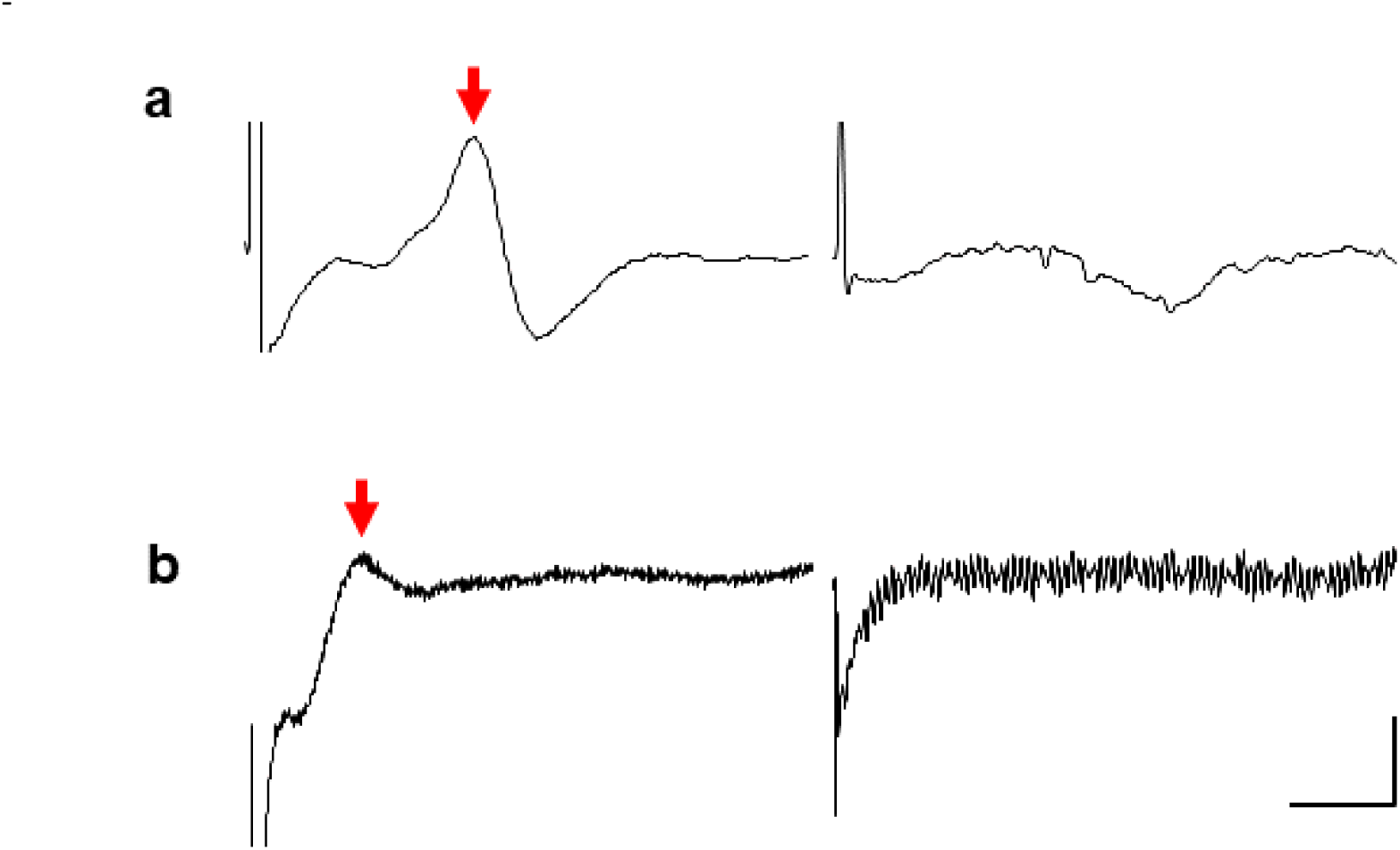
Cerebellar evoked potentials recorded from two patients. **a)** Evoked cerebellar potential (arrow) recorded the right inferior posterior lobe in response to single pulse stimulation of the right tibial nerve from patient S2. The trace on the right shows lack of response from the same recording position following stimulation of the right median nerve. **b)** Evoked cerebellar potential (arrow) recorded from the left posterior lobe in response to a train of stimuli (9 square pulses 0.5 ms, 10 ms interval, 2.7 Hz) delivered to the left forearm. The trace on the right shows no response to the same stimulus train from an adjacent cerebellar recording site. In both examples the trace is an average of 30 consecutive trials. Voltage scale bars = 10µV in **a**, 5µV in **b**; time base =10 ms.

In a second patient (S6), stimulation of the extensor digitorum muscle in the ipsilateral left forearm evoked a response on the surface of the left inferior posterior lobe of the cerebellum (Figure 1b) with an onset latency of approximately 11 ms (time interval between last stimulus in train and onset of response). The evoked potential was confined to one recording site; no responses were found at adjacent recording sites in response to the same stimulus parameters applied to the right forearm. In the remaining five cases (S1, S3-5, S7, 8) no detectable cerebellar responses could be found. In four cases (S4, 5, 7, 8) the absence of a cerebellar response occurred despite the peripheral stimulation evoking a peripheral nerve volley and an SEP recorded over the contralateral parietal lobe (Figure 2). In these cases, it therefore seems reasonable to conclude that the absence of any detectable cerebellar responses was not due to the peripheral stimulation being ineffective in activating ascending sensory pathways.

**Figure 2.**
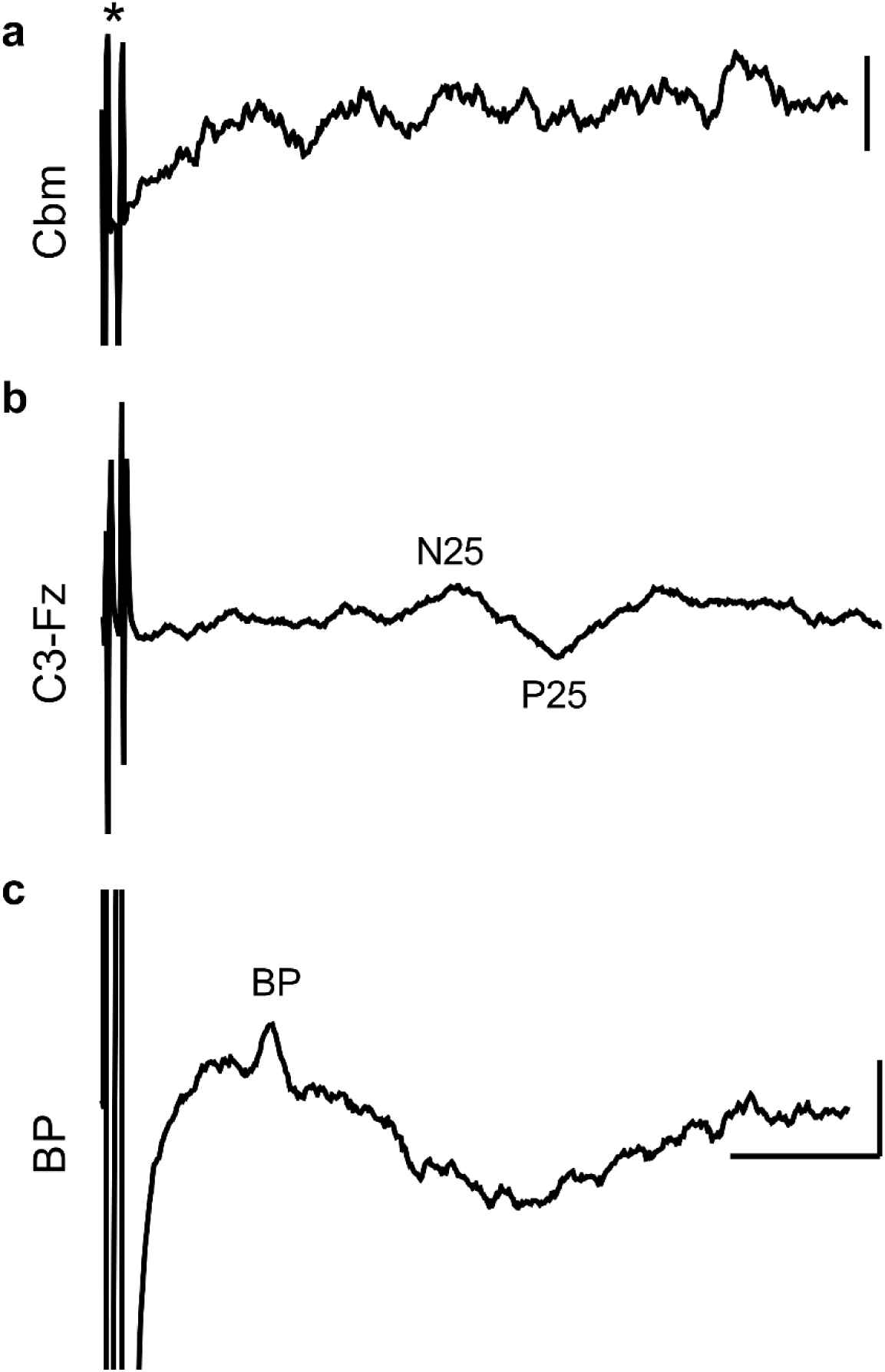
Example results from one patient (S4) with stimulation of the median nerve. **a)** Stimulus artefact indicated by (*) but no detectable cerebellar (Cbm) response was evident. Average of 30 trials. **b)** Cerebral SEPs (N25 and P25) recorded with a scalp electrode (Cz) placed posterior to the central midline over the parietal lobe, referenced to the Fz electrode placed over the frontal midline. **c)** Brachial plexus (BP) peripheral nerve response. All SEP traces based on average of 50trials. Voltage scale bars = 10 µV in **a-c** and 20 µV in **d**; time base =10 ms.

### Motor mapping

Cerebellar stimulation was attempted in a further five patients (cases M1-M5). Cerebellar cortical stimulation did not result in any detectable EMG activity in the peripheral muscles recorded (nasalis and orbicularis oris supplied by the facial nerve, biceps, extensor digitorum communis and flexor carpi radialis in the forearm, small hand muscles, quadriceps, tibialis anterior and abductor hallucis). In one patient (M4) pre-operative fMRI mapping of the motor area in the cerebellum was carried out to help localise the area of interest. The BOLD activation in the inferior posterior lobe of the cerebellum was in an area known to represent the motor function of ipsilateral toes (Figure 3 (8)).Despite the use of this additional fMRI information to help guide location of the cerebellar cortical stimulation, no peripheral muscle EMG responses could be found. However, the BOLD activation area was deep within the right cerebellar hemisphere, (approximately 8 mm below the site of surface stimulation) so the stimulation was unlikely to be effective.

**Figure 3.**
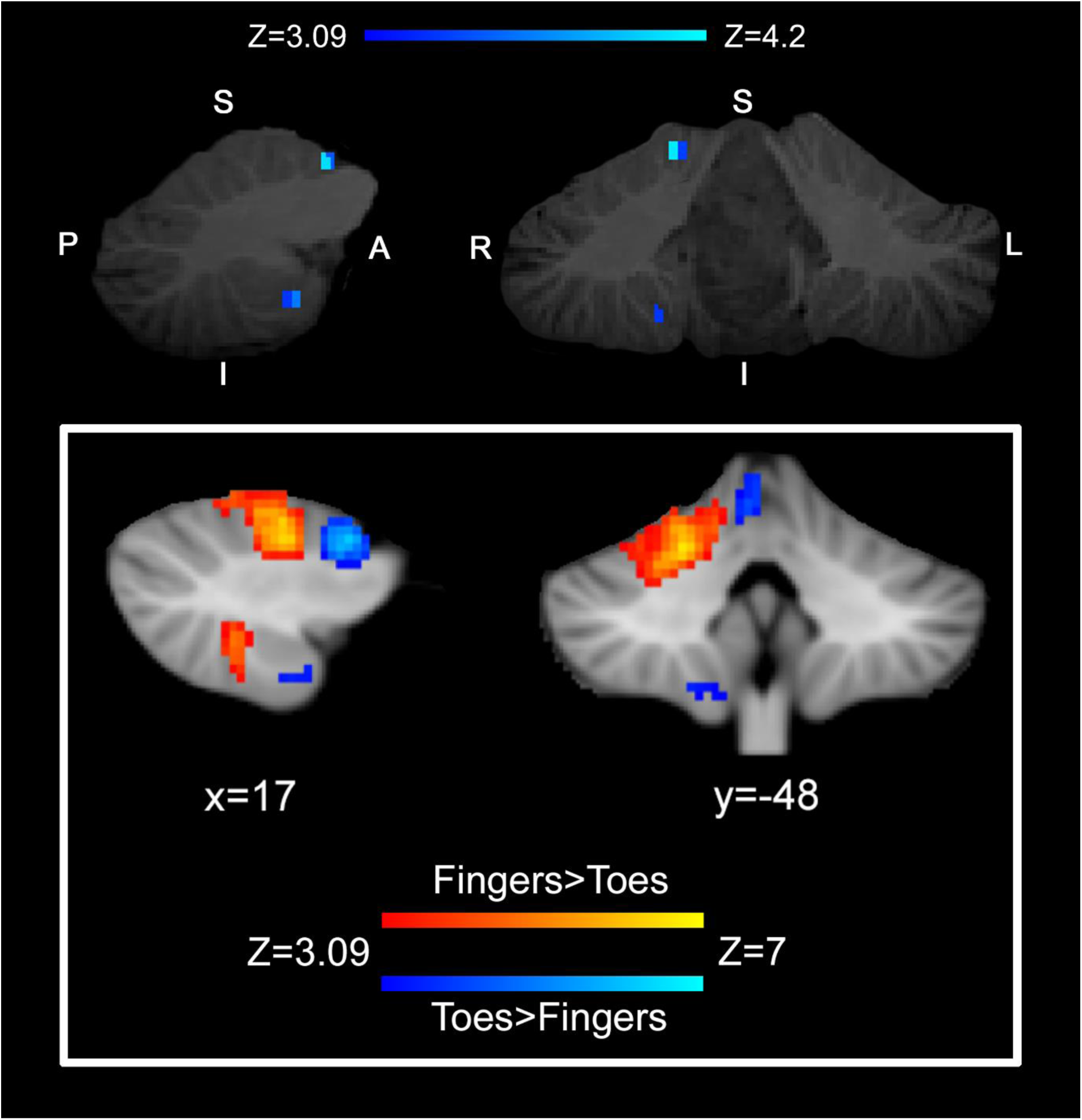
fMRI cerebellar mapping results for a single patient (M4) undergoing posterior fossa ependymoma resection. Top figure demonstrating data representing the contrast between activation from movement of the toes greater than movement of the fingers, shown in blue/light-blue on sagittal and coronal sections, with an uncorrected significance threshold of P<0.001 (i.e. Z=3.09). The reverse contrast did not yield any active areas at the chosen threshold. Note that the activated region in the inferior posterior cerebellar lobe, approximately 8mm from the cerebellar surface. For comparison, bottom inset shows group analysis results (data modified from (8)). demonstrating fingers and toes sensorimotor representation in the cerebellum in 20 healthy participants. The x and y coordinates of group activity are shown in the space of the Montreal Neurological Institute template (in millimetres) - with a cluster forming threshold of Z>3.09 and corrected significance P<0.05. Comparison of the two panels shows good agreement between the patient data and that obtained from the group of 20 healthy controls performing the same tasks.

## Discussion

In this study, we attempted to record evoked field potentials from the exposed surface of the human cerebellum in response to peripheral nerve stimulation in eight patients; and in a further five patients we attempted to record evoked EMG responses in response to direct stimulation of the cerebellar surface. Only peripheral stimulation was successful in evoking responses and this was limited to two out of eight patients (25% of cases). In both successful cases the responses were recorded on the surface of the inferior posterior cerebellar hemisphere, ipsilateral to the site of limb stimulation.

The waveform and onset latencies to upper limb stimulation recorded under anaesthesia in the present study are consistent with those obtained using MEG in awake humans (15, 16). Our findings can also be compared with results obtained from animal studies. For example, in the awake cat, cerebellar cortical responses identified as climbing fibre in origin and evoked by stimulation of the superficial radial nerve in the ipsilateral forelimb can be evoked in the anterior (lobule V) and posterior lobes (lobule VII, rostral paramedian lobule) with an onset latency ranging between 9-14 ms (29, 30). This is similar to the onset latency for climbing fibre responses reported in anaesthetised cats, rats and ferrets (23, 31–33). By comparison, responses in animal experiments attributable to activation of spino-cerebellar pathways terminating as mossy fibres have an onset latency of ~5 ms (34–36) Taken together this suggests that spino-cerebellar pathways can be activated in a range of species by upper limb stimulation. Assuming roughly similar conduction velocities in the ascending tracts and given the longer conduction distances in human, this suggests that the cerebellar responses in human may be mainly mossy fibre in origin.

Consistent with this interpretation are our results from lower limb stimulation. In the present study under anaesthesia, the onset latency of these responses were ~12 ms. By comparison, the onset latencies of climbing fibre responses evoked by hindlimb stimulation and recorded in homologous regions of the posterior lobe of the cerebellum in anaesthetised rats is longer at about 16-19 ms (23). Unless spino-olivocerebellar pathways in human are much faster in conduction than in other species, this suggests that the responses recorded in the present report are mainly mossy fibre in origin, but this would require Purkinje cell recording to verify.

### Methodological considerations

In an attempt to increase the success rate of recording cerebellar evoked responses, a number of changes were made to the peripheral stimulation protocol. This included changes to frequency of stimulation to replicate the parameters used in previous studies. For example, a paired pulse protocol is known to facilitate cerebellar responses (23). And similar peripheral stimulus parameters to those used by Mottolese and colleagues (personal communication) to evoke potentials in the human cerebellum were also attempted. With the caveat of not being able to draw firm conclusions from a small sample size, it was not evident that any of these changes significantly improved our success rate (1 in 4). The finding in one patient that BOLD activation was located deep withing the cerebellum (consistent with a previous fMRI report, (8)); and the effect of anaesthetic on transmission on spino-cerebellar pathways may be important factors to explain our relatively low yield of results, as anaesthetics are known to have a profoundly depressing effect on such pathways (35, 37, 38).

Cerebellar cortical stimulation was also attempted in five additional patients however, no EMG responses were found. As with the sensory stimulation, we tested various stimulation parameters previously used in animal studies (39), in addition to standard parameters used for transcranial MEP monitoring but without success. This negative finding contrasts somewhat with those of Mottolese et al., (17) who reported evoked EMG responses in humans as a result of surface stimulation of the cerebellum. However, a number of factors are likely to have contributed to our negative findings. Chief among these is the report by Mottolese et al., (17) that only 8% of cerebellar stimulation sites evoked a detectable EMG response. They also used a biphasic stimulation protocol which may be critical by increasing the overall electric charge delivered to the cerebellum. It is also noteworthy that in a preliminary report, cerebellar stimulation in patients undergoing posterior fossa surgery has been shown to *indirectly* affect muscle activity;cerebellar stimulation reduced the amplitude of transcranial motor evoked potentials recorded from the contralateral upper limb (40). An additional important consideration in the current study, is that most patients had cerebellar disease. The anatomical distortion and damage from tumours made motor and sensory mapping more challenging. For this reason, it was not possible to precisely localise recording or stimulation sites according to cerebellar anatomy.

## Conclusion

The present study demonstrates it is possible to record under anaesthesia responses from the surface of the human cerebellum evoked by peripheral stimulation. However, a more extensive study would be required to optimize stimulation and recording parameters before such an approach could be used intraoperatively to reliably monitor cerebellar somatosensory function.

## Acknowledgements

This work was supported by Action Medical Research SP4619 (R.A., N.L.C., R.J.E.). We would like to thank Rachel Bissett for her help with the figures.

## Compliance with Ethical Standards

This study was in compliant with Ethical Standards.

